# Why is there so much variability in crop multi-model studies?

**DOI:** 10.1101/2023.02.03.526931

**Authors:** Daniel Wallach, Taru Palosuo, Henrike Mielenz, Samuel Buis, Peter Thorburn, Senthold Asseng, Benjamin Dumont, Roberto Ferrise, Sebastian Gayler, Afshin Ghahramani, Matthew Tom Harrison, Zvi Hochman, Gerrit Hoogenboom, Mingxia Huang, Qi Jing, Eric Justes, Kurt Christian Kersebaum, Marie Launay, Elisabet Lewan, Ke Liu, Qunying Luo, Fasil Mequanint, Claas Nendel, Gloria Padovan, Jørgen Eivind Olesen, Johannes Wilhelmus Maria Pullens, Budong Qian, Diana-Maria Seserman, Vakhtang Shelia, Amir Souissi, Xenia Specka, Jing Wang, Tobias K.D. Weber, Lutz Weihermüller, Sabine J. Seidel

## Abstract

It has become common to compare crop model results in multi-model simulation experiments. In general, one observes a large variability in such studies, which reduces the confidence one can have in such models. It is important to understand the causes of this variability as a first step toward reducing it. For a given data set, the variability in a multi-model study can arise from uncertainty in structure or in parameter values for a given structure. Previous studies have made assumptions about the origin of parameter uncertainty, and then quantified its contribution, generally finding that parameter uncertainty is less important than structure uncertainty. However, those studies do not take account of the full parameter variability in multi-model studies. Here we propose estimating parameter uncertainty based on open-call multi-model ensembles where the same structure is used by more than one modeling group. The variability in such a case is due to the full variability of parameters among modeling groups. Then structure and parameter contributions can be estimated sing random effects analysis of variance. Based on three multi-model studies for simulating wheat phenology, it is found that the contribution of parameter uncertainty to total uncertainty is, on average, more than twice as large as the uncertainty from structure. A second estimate, based on a comparison of two different calibration approaches for multiple models leads to a very similar result. We conclude that improvement of crop models requires as much attention to parameters as to model structure.

## 1. Introduction

Process-based crop models are widely used tools in agronomy, to help analyze past results and to explore scenarios related to alternative management options, effect of climate change, impact of new varieties etc. (Boote et al., 2010). Since the inception of the Agricultural Modeling Inter-comparison and Improvement Project (AgMIP, Rosenzweig et al., 2013), it has become common to organize multi-model simulation studies, where multiple modeling groups are provided with the same input data (i.e. weather, crop management, soil characteristics and initial conditions) and the same observational data for model calibration, and simulate the same response variables. Such studies have shown that the variability in simulated values is very substantial (Asseng et al., 2013; Durand et al., 2018; Salo et al., 2016; Sándor et al., 2020; Wallach et al., 2021a, 2021b). In several studies where multiple crop models were combined with multiple global circulation models (GCMs), it was found that the uncertainty contribution from crop models is as large or larger than that from the GCMs (Asseng et al., 2013; Müller et al., 2021; Wang et al., 2024, 2020). The variability in multi-model studies is probably the most pertinent measure of overall crop model uncertainty, since it reflects the current variability in simulated results among modeling groups. The large uncertainty clearly limits the confidence that one can have in the results of crop models, since it implies that results can vary widely depending on the specific model and modeling group providing the results. A reduction in uncertainty from crop models is necessary in order to increase confidence in the results of these models (Maiorano et al., 2017; Rötter et al., 2011).

In multi-model studies, all modeling groups are given the same input and observation data. Thus the variability between groups has two possible origins, model structure and model parameters. Crop model structure encompasses both the processes taken into account in the models and the specific forms of the equations used to model those processes. The second source of variability is uncertainty in the parameter values. Different groups using the same model structure can have different values for the parameters (Albanito et al., 2022; Confalonieri et al., 2016), again leading to differences in simulated values. It is important to quantify the contributions of structure and parameter uncertainty to overall uncertainty, as a first step in reducing the overall uncertainty. In general, uncertainty is summarized as a variance value, so the question is the relative contributions of structure and parameter variance to the overall variance of simulated values in multi-model studies.

To proceed, it is helpful to have a clear definition of the parameter and structure uncertainties of interest. We propose that an appropriate definition of model structure uncertainty is the range of plausible crop model structures, where by “plausible” we mean a structure that would be deemed acceptable by researchers knowledgeable in the field of crop modeling. The structures represented in open-call multi-model studies can be considered a random sample of plausible structures; random because all modeling groups were invited to participate, plausible because the structures are used by researchers in the field. Our definition of parameter uncertainty for a given model structure and data set is the range of plausible parameter vectors for that structure and data. Here “plausible” means that research groups working with that structure would consider those parameters, or the way in which the parameters are obtained, reasonable. Note that parameter uncertainty is nested within structure uncertainty. That is, the uncertainty in question is different for each model structure, since each structure has its own parameterization. Overall parameter uncertainty is an average over the uncertainties for different structures. Parameter uncertainty covers both the plausible range of parameter default values and the plausible range of calibration procedures, including the choice of which parameters to fit to the data and the procedures of doing the fitting. Each parameter vector represented in open-call multi-model studies is plausible, since it is derived by a practicing research group. For a given model structure, each parameter vector can be considered a random draw from among plausible vectors for that structure, that is, all plausible parameter vectors have an equal chance of being represented in the multi-model study.

A major source of parameter uncertainty is variability in calibration approach. Calibration of crop models and more generally process based models in other fields is difficult because of the large number of parameters and the multiple types of observed variables. As a result, there is no general consensus on the best calibration approach for crop modes (Ahuja and Ma, 2011). One question is how to choose which parameters to estimate and which to treat as fixed. Various studies have proposed sensitivity analysis approaches for choosing the parameters to estimate (e.g. Zadeh et al., 2022 for water quality modeling; Zhang et al., 2014 for rice phenology). Other studies have compared different algorithms for searching for the best parameter values (e.g. Gao et al., 2020; Harrison et al., 2019). The choice of objective function is also variable, and several choices are possible (e.g. Houska et al., 2015 for ecological models). Wallach et al. (2021c) specifically studied the range of calibration approaches used by different modeling groups in three multi-model studies. They found that some groups used a frequentist approach, with various objective functions, and others a Bayesian approach, with various assumptions about the likelihood function. There were also differences in the way to choose the parameters to estimate, with some groups basing the choice on sensitivity analysis, others on expert knowledge about the most important model parameters, sometimes combined with trial and error to choose the parameter set that gives the best fit to the data. Even for modeling groups using the same model structure, there were multiple differences in calibration approach, including different choices of which parameters to fit to the data.. Albanito et al. (2022) also emphasized that different modeling groups make different decisions about calibration practices. In order to quantify the variability in plausible parameter values, we need to take into account all these differences in calibration approach.

Several previous studies have specifically quantified the contributions of structure and parameter uncertainty to total uncertainty for crop models or other process-based models. In these studies, a single modeling group generates multiple parameter vectors for each of several models. Zhang et al. (2017) considered five different models used to predict rice phenology. For each model, 50 parameter sets were generated, based on an assumed uncertainty range for each parameter. Analysis of variance was used to estimate the separate contributions of structure and parameter uncertainty. Tao et al. (2018) used a similar approach for an ensemble of seven models for simulating barley growth and yield. Parameter ranges were based on expert opinion. Both studies identify a fixed subset of parameters that are considered uncertain. These approaches are appropriate for estimating *a priori* uncertainty in a fixed subset of parameter, when no data is available for calibration. However, in multi-model studies, and in our definition of plausible parameter values, models are calibrated using provided data. Calibration modifies the uncertainty in parameter values compared to their *a priori* range. Furthermore, these studies ignore uncertainty in the choice of parameters to estimate. In practice, different modeling groups using the same model structure choose to estimate different parameters, but this variability is not taken into account in these studies. ed. Thus, this approach does not provide a sample of plausible parameter values according to our definition, and is therefore not appropriate for estimating the contribution of parameter uncertainty to total uncertainty in multi-model studies.

A different approach assumes that parameter uncertainty derives from error in observations. The variance of observation error is estimated and then used to estimate the distribution of the parameter estimators. Thus Wallach et al (2017) estimated parameters of two models using the standard regression technique of generalized least squares, which also provides estimates of the variance-covariance matrix of the parameter estimators (Seber and Wild, 1989). Then samples of parameter vectors were generated for each model, and analysis of variance was used to estimate structure and parameter variance. Another approach explores the parameter space around the optimal parameter values, and accepts parameter values that satisfy a goodness of fit test, which is a way of relating parameter uncertainty to observation uncertainty. Migliavacca et al. (2012), looking at 12 budburst models, used this approach to generate 1000 parameter vectors that give fits to the data that are statistically equivalent to the fit using the best parameter values for each model. However, while the effect of observation error on parameter uncertainty is important, it is not relevant to uncertainty in multi-model studies, which use a fixed data set. The variability in parameter values between different modeling groups in multi-model studies is not due to observation error, since all groups use exactly the same data. The variability between groups using the same model structure is due to differences in the choice of default parameter values and in calibration approach, which is not captured in these studies.

In a somewhat different context, Smallman et al. (2021) studied structure and parameter uncertainty for five terrestrial ecosystem models. Bayesian calibration was used to determine parameter distributions for each model. This approach evaluates parameter uncertainty as the result of *a priori* uncertainty and observation error. Since as noted above observation error is not considered in open-call multi-model ensembles, this approach again does not recreate the variability of plausible parameter values, according to our definition.

Xiong et al. (2020) considered structure and parameter contributions to uncertainty in wheat model simulations applied to two Chinese regions. Three models were considered. Parameter variability was due to different levels of spatial detail (same parameters at all locations, different parameters for each major agro-ecological zone, parameters fit to phenology data or parameters fit to both phenology and yield data). In this case parameter uncertainty measures the variability between different specific choices of the way to determine the parameter values. The approach is not aimed at reproducing the variability in multi-model studies.

These previous approaches have sampled from specific types of parameter uncertainty, namely a priori uncertainty, uncertainty due to uncertainty in the data, or uncertainty related to different choices of the way to treat spatial data. None of these approaches is designed to recreate the parameter variability between different groups using the same structure in multi-model studies. That variability results from differences in default parameter values and from multiple differences in calibration approach. To date, there do not seem to have been any studies which realistically take into account the full uncertainty in plausible parameter values.

The purpose of this study is to propose, for the first time, estimates of structure and parameter uncertainty in multi-model studies that fully account for parameter uncertainty. Our case study concerns prediction of wheat phenology. Crop phenology is important since it has a major effect on crop yield and is a major aspect of crop response to global warming (Ahmad et al., 2019; Fatima et al., 2020; Liu et al., 2022). Process-based models of phenology, usually embedded in crop or agroecosystem models, are often used to predict the effect of weather on phenology (Bindi et al., 2015; Muleke et al., 2022). Multi-model studies specifically of crop phenology have shown wide variability in simulated results (Wallach et al., 2021b, 2021a). Thus, the uncertainty in prediction of crop phenology is of interest in its own right, as well as serving as an example for crop models more generally.

We estimate parameter uncertainty based on the differences between different modeling groups using the same model structure, in multi-model studies. This automatically takes into account all the differences in choice of default parameters and in calibration approach between different groups.Specifically, we revisit three published open-call multi-model studies on predicting wheat phenology (Wallach et al., 2021b, 2021a), where some model structures were used by more than one modeling group. Given the results from multiple different model structures, and from multiple groups using the same structure, we estimate structure and parameter variances using random effects analysis of variance. Our approach, therefore, provides more realistic estimates for the relative contributions of structure and parameter uncertainty than have previously been available The methodology employed here, namely using random effects analysis of variance on data from multi-model studies, is applicable in general to process-based models.

In the particular cases treated here, we can obtain a second estimate of the variance in simulated values due to parameter uncertainty. In the multi-model studies revisited here, each participating group used their “usual” calibration approach. Subsequently an original calibration protocol was developed, and most of the groups redid the calibration using the new protocol. Thus for multiple groups we have results for two different calibration procedures, “usual” and “protocol”, both of which may be considered plausible The comparison between them can be used to estimate potential variability in parameters due to calibration approach. These results can be compared with the results based on the original multi-model studies.

## 2. Materials and Methods

### 2.1 Observational data

The multi-model studies use three different data sets (**Table 1**). Two of the data sets are from winter wheat variety trials in France. These two data sets have identical structures (same sites and sowing dates) but concern two different wheat varieties, “Apache” and “Bermude”. Both varieties are of intermediate precocity, but Bermude has a longer cycle. The observed data from each environment consist of dates of stem elongation (BBCH30 on the BBCH scale, Meier, 1997) and of the middle of heading (BBCH55). The third data set is from a multi-site sowing date experiment on spring wheat variety Janz in Australia. The original data were observations of growth stage at weekly intervals. The data provided to modeling groups were the dates of each BBCH stage, from the first to the last observed stage, derived from interpolation of the original data. All data sets were split into two subsets, one for calibration and the other for evaluation. The calibration and evaluation data sets had neither site nor year in common. Thus evaluation was a true test of how well models could predict out-of-sample conditions.

**Table 1.** Data sets. **The three data sets used in the multi-model ensemble studies analyzed here. The French data sets are described in** (Wallach et al., 2021a). **The Australian data set is described in** (Wallach et al., 2021b).

| Data set | Number of calibration environments | Number of evaluation environments | Observed development stages* | Simulated development stages used for analysis* |
| --- | --- | --- | --- | --- |
| France<br>Variety Apache | 14 | 8 | BBCH30, BBCH55 | BBCH10, BBCH30, BBCH55 |
| France<br>Variety Bermude | 14 | 8 | BBCH30, BBCH55 | BBCH10, BBCH30, BBCH55 |
| Australia<br>Variety Janz | 24 | 18 | BBCH stage at weekly intervals | BBCH10, BBCH30, BBCH65, BBCH90 |

\*The development stages, using the BBCH scale (Meier, 1997) are emergence (BBCH10), start of stem elongation (BBCH30), middle of heading (BBCH55), full flowering (BBCH65), and maturity (BBCH90).

### 2.2 Participating modeling groups and model structures

The call for participants in these studies was published using the mailing lists of AgMIP and of several crop models. All modeling groups that volunteered (a modeling group is one or more individuals who work together to run a model) were accepted. No specific attempt was made to encourage or discourage participation by different groups that used the same structure. In the event, there were two or three (depending on the simulation study) model structures used by more than one group (Table 2). These are thus what we have termed “open-call” multi-model studies.

**Table 2.** The three open-call multi-model simulation studies analyzed here (Wallach et al., 2021a, 2021b).

| Simulation study | Data set (country, cultivar) | Calibration procedure | Number of modeling groups | Repeated structures* |
| --- | --- | --- | --- | --- |
| multi-model study 1 | France, Apache | Usual | 27 | S2 (4), S4 (3), S18 (2) |
| multi-model study 2 | France, Bermude | Usual | 27 | S2 (4), S4 (3), S18 (2) |
| multi-model study 3 | Australia, Janz | Usual | 28 | S2 (3), S4 (3), S18 (2) |
| multi-model study 4 | All 3 data sets above | Protocol | 19 | not relevant |

\*The model structures used by more than one modeling group, and in parentheses, the number of groups that used that structure.

A total of 30 modeling groups using 23 different structures participated in one or more of the simulation experiments. For the purposes of this study, each model with a unique name was considered as a different model structure. A list of structures represented here, together with references that describe the model equations, is given in Supplementary Table 1. In all cases, the phenology models were embedded in general crop models, so the model structure here refers to the overall crop model. Most but not all of the groups participated in all the studies.

In the original reports of these studies, the different modeling groups were identified by a code (M1, M2 etc.) without identifying the group or the model structure used (Wallach et al., 2021b, 2021a). The same codes are used here. In addition, we assign here codes to the different model structures (S1, S2, etc.). The same model structure can be associated with more than one modeling group.

### 2.3 Simulation studies

The multi-model simulation studies, based on the three data sets, are summarized in Table 2. In the first two studies, based on the two French data sets, each participating modeling group implemented their usual calibration method to calibrate their model using the calibration data subsets and then simulated days from sowing to development stages BBCH10, BBCH30, and BBCH55 for both the calibration and evaluation environments (Wallach et al., 2021a) (Table 2). In the study based on the Australian data set (Wallach et al., 2021b), each participating modeling group again used their usual calibration method and then simulated days from sowing to development stages BBCH10, BBCH30, BBCH65, and BBCH90 for the Australian environments. Neither the French nor the Australian data sets had observations of BBCH10 (emergence), which was included in the variables to simulate in order to have an example of a simulated stage without calibration data. Detailed information about the variability in calibration procedures among the different modeling groups is presented in Wallach et al. (2021c).

Following the three multi-model studies, a standardized calibration procedure (hereafter the calibration “protocol”) was proposed (Wallach et al., 2023). In a fourth simulation experiment, each of 19 participating groups used the protocol calibration to simulate for the same three data sets used in the first studies. Any group that participated in the fourth study but that had not participated in the first three studies first did the calibration using their usual method, before applying the protocol. Thus for each data set and each participant, there were two parameter vectors, based on usual or protocol calibration.

### 2.4 Estimation of structure and parameter variance

#### Variance components approach

We assume that for each of the open-call simulation experiments, the model structures are a random sample from plausible structures. We further assume that each parameter vector is a random draw from plausible parameter vectors for the corresponding model structure. The statistical model for a simulated value is

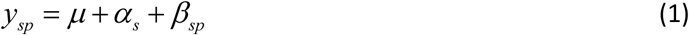

where *y_sp_* is the simulation result for the output y (for example days to BBCH30 in a particular environment) obtained using the s^th^ draw of model structure *s*and the p^th^ draw of parameter vector for that model structure. µ is the overall mean of the simulation results, *a_s_* is the effect of model structure and *β_sp_* is the effect of parameter vector nested within the model structure. The parameter effect is nested within structure, because different structures have different parameterizations. There is no residual term because these are simulated values with no measurement error.

In a fixed effects analysis of variance, we are mainly interested in the effects *α* and *β* of the specific structures and parameters of the sample. Here, on the other hand, we are mainly interested in the variability between model structures and between parameter vectors for each given structure. The appropriate analysis tool is a random effects analysis of variance.

In random effects analysis of variance, *a_s_* and β*_sp_* are treated as random effects. *a_s_* is random because the structure chosen at the s^th^ draw could be any plausible structure. All the *a_s_* have the same distribution, because every draw is from the same population of model structures. We are interested in the variance of the *a_s_*, noted 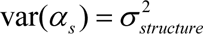 for all s. Similarly, each β*_sp_* has the same distribution. We note 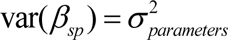 for all *sp*. The random terms are assumed independent so that the total variance for one environment and one simulated variable is the sum of model structure variance and parameter variance (Scheffé, 1959):

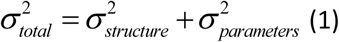

The fraction of total variance due to parameter variance is then 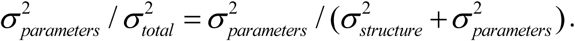

The variances 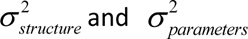 can be estimated using random effects analysis of variance. In the case of a balanced sample (same number of parameter vectors for each model structure), there are analytical expressions for the maximum likelihood (ML) estimators of the variances. In the general, unbalanced case, the estimators are calculated iteratively. We use the lmer function of the lme4 package (Bates et al., 2015) in R (R Core Team, 2017), which is designed to analyze linear mixed effect models. We use the REML option, which corrects for the fact that the ML estimators are biased. Commented R script for calling the lmer function and for extracting the variances of interest are shown in Supplementary.

An illustration of the data used in this approach is given in Fig. 1. As that figure shows, three of the model structures (S2, S4, and S18) were used by more than one modeling group. It is the variability between the different groups using the same model structure that allows us to estimate parameter variance. The results for all groups contribute to the estimation of structure variance. Note that the simplified approach to estimating parameter uncertainty using these same data in Wallach et al. (2021a, 2021b) underestimates the contribution of parameter uncertainty.

**Figure 1.**
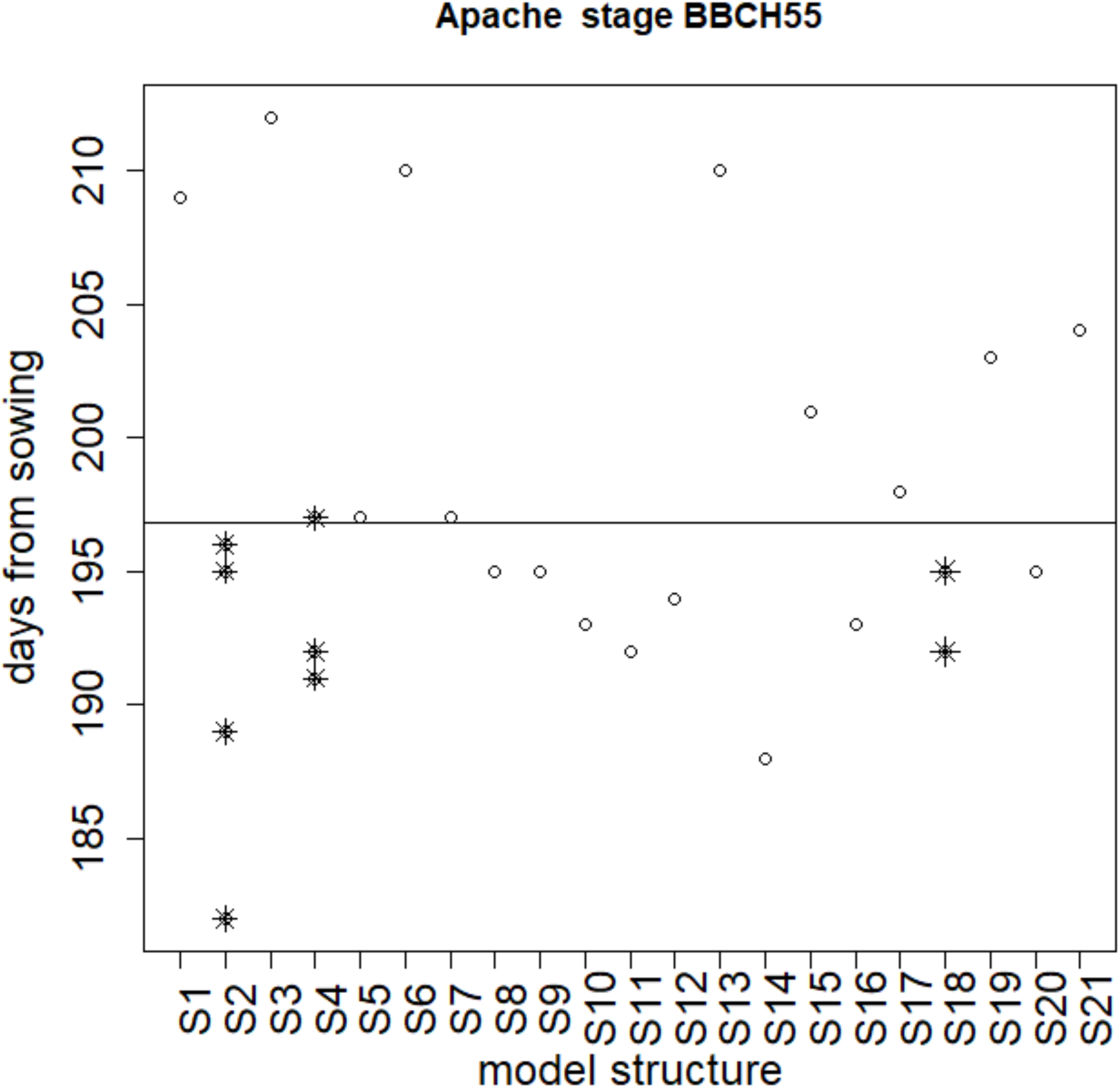
Example of data used for estimating parameter and structure uncertainty in the “variance components” approach (location Foreste, sown 10/11/2011). Each circle or star is a simulated value of days from sowing to mid-heading (stage BBCH55) for variety Apache in one environment from simulation study 1. All modeling groups used the same inputs. The x-axis shows the model structure. In most cases, each structure was used by a single modeling group (open circles), but structures S2, S4, and S18 were used respectively by 4, 3, and 2 groups (data shown as stars). It is the variability between simulations from different groups using those model structures that is the basis for estimating parameter uncertainty. The horizontal line is the average of all simulated values.

**Figure 2.**
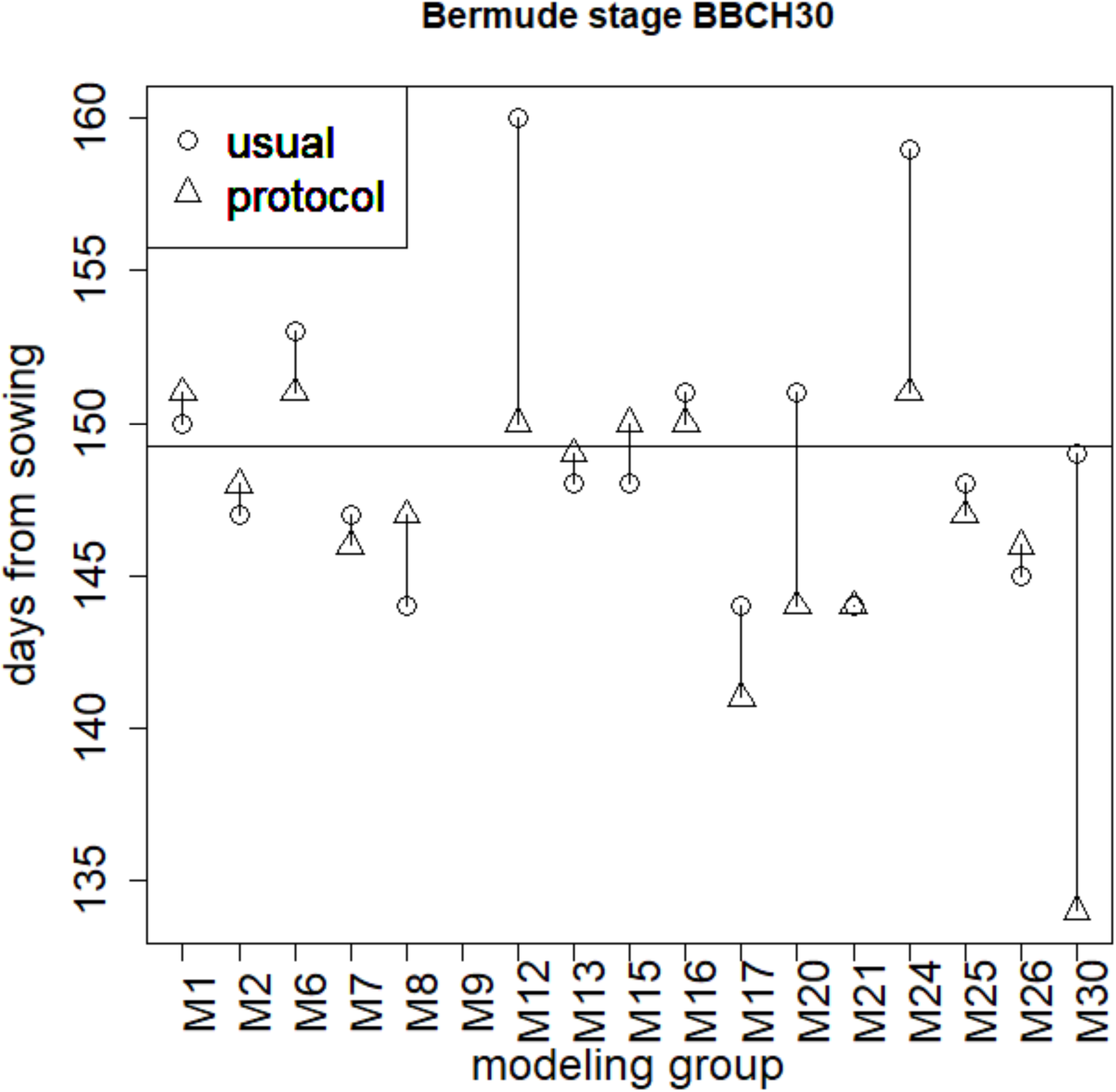
Example of simulated values where each group used two calibration approaches. Simulations are of days from sowing to stem elongation (stage BBCH30) for variety Bermude, location Boigneville, sown 21/10.2013. Each code (M1, etc.) refers to a specific modeling group (x-axis). For each modeling group, there are two simulated values, one using parameters based on usual calibration and the other using parameters based on protocol calibration. The differences between the two values are used to estimate parameter variance. The horizontal line shows the mean of simulations using usual calibration.

#### Comparison of usual and protocol calibration

A second estimation of parameter uncertainty is based on comparing the results between usual and protocol calibration, for each group. The major difference between the calibration approaches was in the choice of which parameters to estimate from the data. For “usual” calibration, there was a wide diversity of methods of choosing parameters to estimate (Wallach et al., 2021c). None of the calibration methods was explicitly designed to avoid over-fitting. The new protocol based the choice of parameters to estimate on standard statistical model selection methods. For each development stage in the calibration data, one parameter that had a similar effect in all environments was automatically chosen to be calibrated. This was usually the number of degree days to the stage. Then additional “candidate” parameters were identified. These were tested in a procedure like forward regression. A candidate was chosen to be estimated if it led to a reduction in the corrected Akaike Information Criterion (AICc). This criterion has the form of a penalized likelihood; it has a term which decreases if the error in fitting the data decreases, and a term which increases as the number of estimated parameters increases. It is designed to avoid-over-fitting.

For each environment, we calculate the variance of the difference between the two calibration approaches (usual and protocol), noted var*_i_* for modeling group *i* (eq. 1), and then average over modeling groups to obtain an estimated parameter variance for that environment (eq. 2).

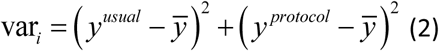

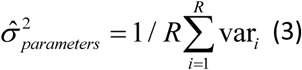

where 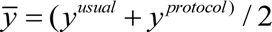 and R is the number of modeling groups.

We do not have a rigorous method of estimating structure variance from these data. The difficulty is that the assumptions underlying the random effects model described above are not satisfied; the parameter vectors obtained by protocol calibration are not independent between modeling groups, since all use the same protocol. Therefore, or each environment we use the estimate of structure variance from the variance components approach above.

## 3. Results

Initially, all calculations were done for the calibration and evaluation data subsets separately. However, there were no systematic differences between the two (Fig. S1 and Fig. S2). Since both calibration and evaluation data are sampled from the same target population, this is perhaps not surprising. All results here are therefore based on the combined calibration and evaluation data. Table 3 and Table 4 show parameter variance, and parameter variance as a fraction of total variance, averaged over all environments for the French and Australian data sets, respectively. Note that the estimated contribution of parameter variance to total variance is always much less for BBCH10, for which there were no observed values, than for the other simulated development stages.

**Table 3.**
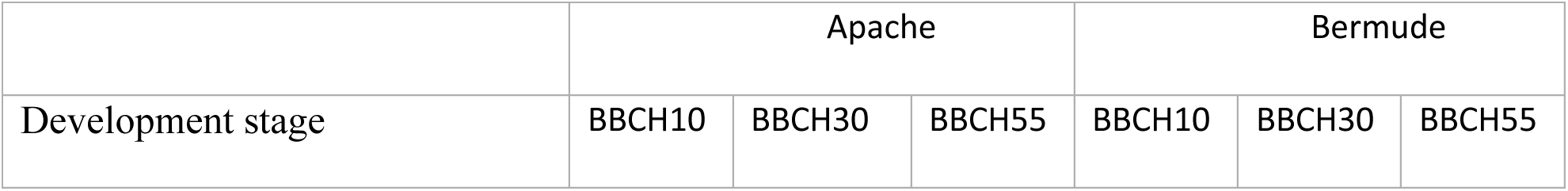
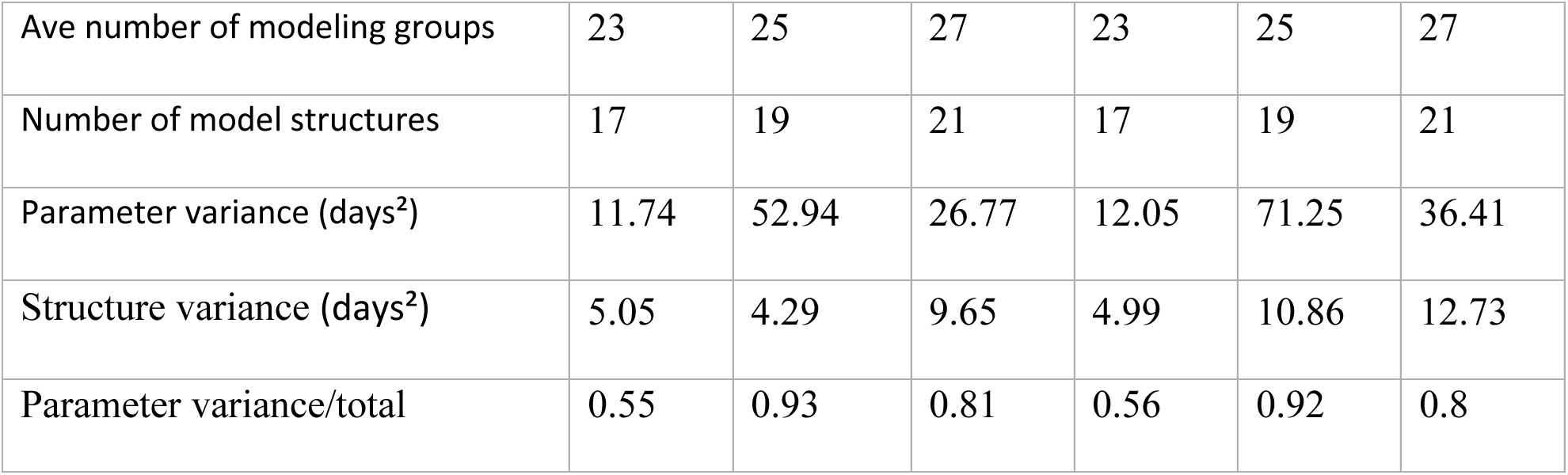
Parameter and structure variance for French data sets estimated using the variance components calculation. The variance components estimate is based on ensemble studies where some model structures were used by multiple modeling groups. Results are averages over environments. The ratio of parameter variance to total (last row) is an average over the ratios for each environment.

**Table 4.** Parameter and structure variance for the Australian data set estimated using the variance components calculation. The variance components estimate is based on ensemble studies where some model structures were used by multiple modeling groups. Results are averages over environments. The ratio of parameter variance to total (last row) is an average over the ratios for each environment.

| Development stage | BBCH10 | BBCH3<br>0 | BBCH65 | BBCH90 |
| --- | --- | --- | --- | --- |
| Average number of modeling groups | 23 | 24 | 28 | 26 |
| Average number of model structures | 17.95 | 19.93 | 22.64 | 21.05 |
| Parameter variance (days<sup>2</sup>) | 40.69 | 102.88 | 47.1 | 30.3 |
| Structure variance (days<sup>2</sup>) | 276.33 | 45.74 | 48.4 | 90.71 |
| Parameter variance / total | 0.12 | 0.66 | 0.49 | 0.24 |

Table 5 shows results averaged over all environments of both the French and Australian data sets, for both the variance-components approach and the approach based on comparing usual and protocol calibration. Separate values are given for BBCH10 and for all other stages. For development stages other than BBCH10, the estimated contribution of parameter uncertainty is large. It ranges from 24 to 93%, and is larger than 50% for 11 out of the 14 combinations of data set, development stage and approach for estimating the contribution of parameter variance.

**Table 5.**
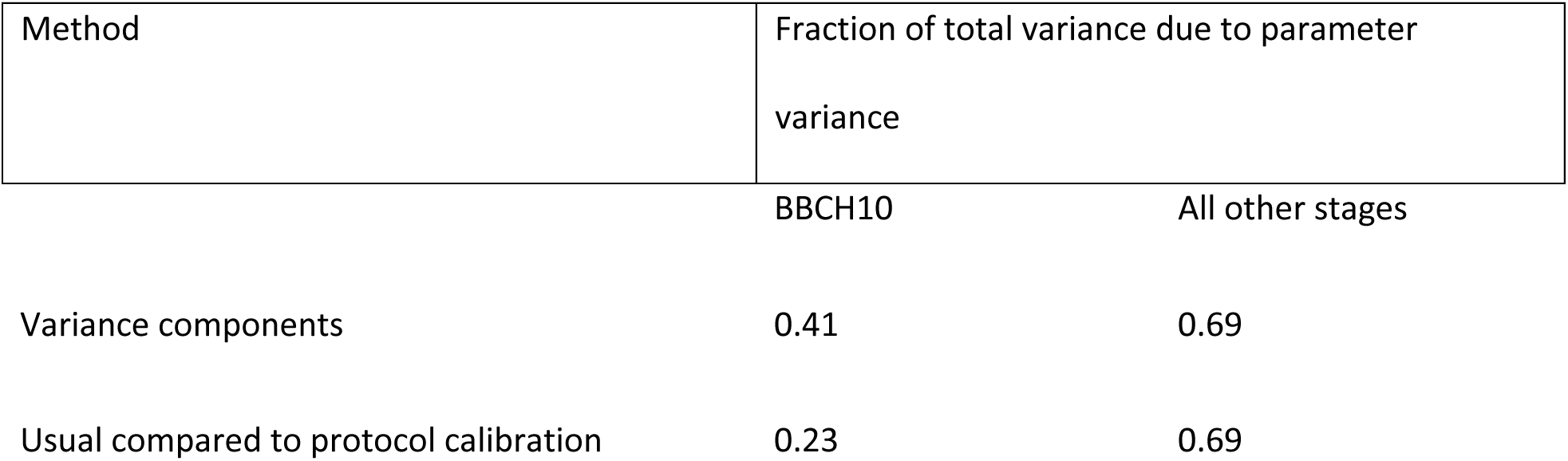
Fraction of total variance due to parameter variance based on two estimation procedures: variance components and comparison of usual and protocol calibration. Averages are over all environments of both the French and Australian data sets, either for the development stage BBCH10 or for all other development stages.

| Method | Fraction of total variance due to parameter variance |  |
| --- | --- | --- |
|  | BBCH10 | All other stages |
| Variance components | 0.41 | 0.69 |
| Usual compared to protocol calibration | 0.23 | 0.69 |

## 4. Discussion

The variability in parameters between different groups using the same model structure results from differences in fixed parameter values and in calibration approach. None of the data sets here included time to emergence (BBCH10), so variability in simulating time to emergence likely depends only on variability in default values. On the other hand, there are data for subsequent development stages, so simulation of those stages will depend on the values of calibrated parameters. The parameter uncertainty contribution to variability in days to emergence is always smaller (41% on average) than the contribution to variability in days to other stages (69% on average). This shows that the importance of parameter uncertainty depends on both the specific variable simulated and on the data available for calibration. It seems that variability in simulated variables affected by calibration is larger than variability in simulated variables which depend just on default parameter values.

When averaging over data sets and simulated variables, for variables other than time to emergence, the resuts show that parameter uncertainty represents 69% of total uncertainty. The same value is obtained using variance components or the comparison of usual and protocol calibration. Structure uncertainty contributes 31%, so the contribution of parameter uncertainty is more than twice as large as the contribution of structure uncertainty. Previous studies that have estimated structure and parameter uncertainty contributions to overall uncertainty of plant or ecosystem models have generally found that structure uncertainty is larger than parameter uncertainty (Tao et al., 2018; Xiong et al., 2020), including studies that have specifically considered simulation of phenology (Migliavacca et al., 2012; Wallach et al., 2017; Zhang et al., 2017). This is in fact not too surprising. Previous studies have quantified the effect of some specific source of parameter uncertainty, such as observation error, or *a priori* uncertainty, or choice of data to use. The study here on the other hand takes into account the full variability in default values and calibration approach between different modeling groups who could participate in open-call multi-model studies.

Several studies have emphasized the uncertainty in parameter values and the necessity of better evaluating parameter uncertainty and improving calibration practices (Iizumi et al., 2009; Ramirez-Villegas et al., 2017; Seidel et al., 2018; Yang et al., 2024). The conclusion from this study, which remains to be confirmed more generally, is that parameter uncertainty, in particular due to uncertainty in calibration approach, is as or more important than structure uncertainty as a cause of variability in multi-model studies. This implies that improving calibration approach, and sharing improved practices widely, should be a major priority of the crop modeling community.

In multi-model study number 4 (Wallach et al., 2023) It was found that total variability between simulated values was reduced by 22% when all groups used the new protocol, compared to the case where each group used its usual calibration approach. Furthermore, prediction error was reduced by 11% by use of the new protocol. This is a clear indication that improved, shared calibration approaches can substantially reduce variability in multi-model studies and thereby enhance confidence in crop model simulations. Recently, building on the calibration protocol for phenology data, a crop model calibration approach has been proposed that is generic (i.e. can be applied to essentially all crop models and data sets). It is directly based on statistical principles, which increases confidence that the approach uses calibration data effectively (Wallach et al., 2025, 2024). This calibration protocol is being tested in several studies, and would be a candidate for a standard calibration approach for crop models.

The present study has several limitations, which must be taken into account before generalizing the results. First, this is a case study, so the specific results refer to the specific cases studied here. Also, there are quite large differences for different environments and development stages (Supplementary Figs. S1, S2. However, the studies use three different data sets, with 22 or 42 different environments each. The French and Australian data sets not only have quite different environmental conditions and plant material (winter wheat in France and spring wheat in Australia), but also quite different measured variables (two development stages for the French data set, weekly development stage for the Australian data set). There were also substantial differences within the French and Australian data sets. For variety Apache, considering just the calibration data, time from sowing to stages BBCH30 and BBCH55 ranged from 126 to 192 days and from 177 to 234 days respectively. For the Australian data, time from sowing to stages BBCH30, BBCH65 and BBCH90 ranged from 49 to 114 days, from 90 to 177 days and from 130 to 205 days respectively. There is substantial variability in results for different environments and different simulated variables, but the range of situations analyzed gives some confidence in the overall conclusion, that parameter uncertainty is a major contribution to overall model uncertainty when simulating phenology. A second limitation is that the study concerns only simulation of phenology. It remains to be seen whether parameter uncertainty is also dominant when simulating crop growth and yield. Calibration in that case is more complex, which may lead to larger calibration uncertainties, but also the simulations involve more equations, which may lead to more structure uncertainty.. Thus it is not clear if the same conclusions as here, that parameter uncertainty contributes more to overall uncertainty than structure uncertainty, also applies in general to crop models In any case, however, it seems probable that taking into account the full variability in plausible parameters will lead to a larger contribution of parameter uncertainty to total uncertainty than previously thought, for all types of process-based models. A third limitation is that the results rely on the variability between different groups that use the same model structure, but only three model structures were used by more than one group, and at most four groups used the same model structure. The results here thus represent only the parameter variability between a small number of groups, using only three models. This is somewhat mitigated by the fact that that we have averaged over a fairly large number of environments, in three different data sets. Most importantly, we also have a second estimate of the contribution of parameter uncertainty, based on the comparison of usual and protocol calibration. This second estimate is based on a sample of two different calibration approaches for all models. The average contribution of parameter to total uncertainty using this second approach, again ignoring simulations of time to emergence, was 69%, essentially the same as the value obtained using the variance components analysis. This increases confidence in the variance component results.

It should also be recognized that the assumption of a random sample of structures and parameters underlying the random effects analysis of variance is probably not exactly satisfied. The modeling groups in this study were not proactively chosen to represent a random sample, but rather were simply the modeling groups that volunteered to participate. Such ensembles have been called “ensembles of opportunity” (Tebaldi and Knutti, 2007). This could for example lead to underestimation of structure uncertainty, if the full range of plausible structures was not sampled. It is not clear how to quantify or correct this. Perhaps the best one can do is to ensure that the invitation to participate is widely disseminated.

The question of the relative importance of different sources of uncertainty in process-based models, and in particular of structure and parameter uncertainty, arises in many fields, such as hydrology (Butts et al., 2004) or earth system models (Ricciuto et al., 2021). Many of these fields also employ multi-model studies (Thébault et al., 2024). The analysis of variance done here can be easily applied to these other fields., in order to estimate the contributions of structure and parameter uncertainty to the overall variability. The requirement is to have the results of an open-call multi-model study where some model structures are used by more than a single modeling group. The analysis proposed here does not require any additional simulations, and the calculations can be easily done using existing software, for example the R software package used here.

## 5. Conclusions

The large variability in crop multi-model simulations reduces the confidence one can have in such simulations, so it is important to understand the causes of this variability. A realistic estimate of parameter uncertainty must take into account the actual variability in parameters between different modeling groups. We illustrate a simple method of doing so, based on results of multi-model studies associated with random effects analysis of variance. This methodology is easily applicable to any case where the necessary multi-model results exist.

The parameter uncertainty taken into account here is different than in previous studies, where specific sources of parameter uncertainty were examined. Unlike those previous studies, we find in the case study here that parameter uncertainty makes the largest contribution to overall uncertainty, substantially larger than structure uncertainty. The larger parameter uncertainty found in the present study is unsurprising, since multiple sources of parameter uncertainty are included here, which suggests that it may be generally true that parameter uncertainty makes a larger contribution to total uncertainty than previously thought.

The importance of parameter uncertainty emphasizes the need for giving more attention to parameter values and in particular to Improved calibration approaches and procedures. Improving parameter values should be a major goal of the crop modeling community, on a par with work on improving model equations.

## Author contribution

Daniel Wallach: Conceptualization, methodology, project administration, writing - original draft, validation. Taru Palosuo, Henrike Mielenz, Peter Thorburn, Samuel Buis, Sabine Seidel: Conceptualization, methodology, project administration, writing, review &editing. All other authors: expertise on models and simulation results, writing, review & editing.

## Supporting information

Supplement_Wallach.etal

## 6. Acknowledgements

This work was in part supported by the Collaborative Research Center 1253 CAMPOS (Project 7: Stochastic Modelling Framework), funded by the German Research Foundation (DFG, Grant Agreement SFB 1253/1 2017), the Academy of Finland through projects AICropPro (316172) and DivCSA (316215), the BonaRes projects ’’Soil3’’ (BOMA 03037514) and “I4S” (031B053I) of the Federal Ministry of Education and Research (BMBF), Germany, the Deutsche Forschungsgemeinschaft (DFG, German Research Foundation) under Germany’s Excellence Strategy - EXC 2070 – 390732324 EXC (PhenoRob), the project BiomassWeb of the GlobeE programme (Grant number: FKZ031A258B) funded by the Federal Ministry of Education and Research (BMBF, Germany), the INRA ACCAF meta-programme, the German Federal Ministry of Education and Research (BMBF) in the framework of the funding measure “Soil as a Sustainable Resource for the Bioeconomy –-BonaRes”, project “BonaRes (Module B): BonaRes Centre for Soil Research, subproject B” (grant 031B0511B), the National Key Research and Development Program of China (2017YFD0300205), the National Science Foundation for Distinguished Young Scholars (31725020), the Priority Academic Program Development of Jiangsu Higher Education Institutions (PAPD), the 111 project (B16026), and China Scholarship Council, the Agriculture and Agri-Food Canada’s Project J-002303 under the Interdepartmental Research Initiative in Agriculture, the DFG Research Unit FOR 1695 ‘Agricultural Landscapes under Global Climate Change – Processes and Feedbacks on a Regional Scale, the U.S. Department of Agriculture National Institute of Food and Agriculture (award no. 2015-68007-23133) and USDA/NIFA HATCH grant N. MCL02368, the National Key Research and Development Program of China (2016YFD0300105), The Broadacre Agriculture Initiative, a research partnership between University of Southern Queensland and the Queensland Department of Agriculture and Fisheries, the Academy of Finland through project AI-CropPro (315896), the JPI FACCE MACSUR2 project, funded by the Italian Ministry for Agricultural, Food, and Forestry Policies (D.M. 24064/7303/15 of 6/Nov/2015), the SYSTEMIC funded by JPI HDHL, JPI-OCEANS and FACCE-JPI under ERA-NET (n.696295), the SustEs project funded by the Ministry of Education, Youth and Sports of the Czech Republic (CZ.02.1.01/0.0/0.0/16_019/000797). The time of MTH and KL was supported by the Grains Research and Development Corporation (Project UOT1906-002RTX). The observed Australian phenology data were jointly funded by CSIRO, the Grains Research and Development Corporation (GRDC) under the “Adding Value to GRDC’s National Variety Trial Network” project (CSA00027). The order in which the donors are listed is arbitrary.

## Notes

### Competing Interest Statement

The authors have declared no competing interest.

### Summary of Updates

The axis titles of Figures 1 and 2 were revised.

