## Supplement_Wallach.etal for "Why is there so much variability in crop multi-model studies?"

Supplementary Information

Wallach^1^, Daniel; Palosuo^2^, Taru; Mielenz^3^, Henrike; Buis^4^, Samuel; Thorburn^5^, Peter; Asseng^6^, Senthold; Dumont^7^, Benjamin; Ferrise^8^, Roberto; Gayler^9^, Sebastian; Ghahramani^10^, Afshin; Harrison^11^, Matthew Tom; Hochman^5^, Zvi; Hoogenboom^12^, Gerrit; Huang^13^, Mingxia; Jing^14^, Qi; Justes^15^, Eric; Kersebaum^16,17,18^, Kurt Christian; Launay^19^, Marie; Lewan^20^, Elisabet; Liu^11^, Ke; Luo^21^, Qunying; Mequanint^9^, Fasil; Nendel^16,17,22^, Claas; Padovan^8^, Gloria; Olesen^23^, Jørgen Eivind; Pullens^23^, Johannes Wilhelmus Maria; Qian^14^, Budong; Seserman^16^, Diana-Maria; Shelia^12^, Vakhtang; Souissi^24^, Amir; Specka^16^, Xenia; Wang^13^, Jing; Weber^25^, Tobias, K.D.; Weihermüller^26^, Lutz, Seidel, Sabine J.*^1^

^1^Institute of Crop Science and Resource Conservation, University of Bonn, Germany

^2^Natural Resources Institute Finland (Luke), Helsinki, Finland

^3^Julius Kühn Institute (JKI) – Federal Research Centre for Cultivated Plants, Institute for Crop and Soil Science , Braunschweig, Germany

^4^INRAE, UMR 1114 EMMAH, Avignon, France

^5^CSIRO Agriculture and Food, Brisbane, Queensland, Australia

^6^Technical University of Munich, Department of Life Science Engineering, Digital Agriculture, HEF World Agricultural Systems Center, Freising, Germany

^7^Plant Sciences & TERRA Teaching and Research Centre, Gembloux Agro-Bio Tech, University of Liege,

Gembloux, Belgium

^8^Department of Agriculture, Food, Environment and Forestry (DAGRI), University of Florence, Italy

^9^Institute of Soil Science and Land Evaluation, Biogeophysics, University of Hohenheim, Stuttgart, Germany

^10^Centre for Sustainable Agricultural Systems, Institute for Life Sciences and the Environment, University of Southern Queensland, Toowoomba, Queensland, Australia

^11^Tasmanian Institute of Agriculture, University of Tasmania, Launceston, Tasmania, Australia, 7248

^12^Agricultural and Biological Engineering Department, University of Florida, Gainesville, Florida, USA & Global Food Systems Institute, University of Florida, Gainesville, Florida, USA

^13^College of Resources and Environmental Sciences, China Agricultural University, Beijing, China

^14^Ottawa Research and Development Centre, Agriculture and Agri-Food Canada, Ottawa, Canada

^15^CIRAD, Univ Montpellier, Persyst Department, Montpellier, France

^16^Leibniz Centre for Agricultural Landscape Research (ZALF), Müncheberg, Germany

^17^Global Change Research Institute CAS, Brno, Czech Republic

^18^Institute for Tropical Plant Production and Agricultural Systems Modelling, Georg-August University, Göttingen, Germany

^19^INRAE, US 1116 AgroClim, Avignon, France

^20^Department of Soil and Environment, Swedish University of Agricultural Sciences (SLU), Uppsala, Sweden

^21^Hillridge Technology Pty Ltd, Sydney, Australia

^22^Institute of Biochemistry and Biology, University of Potsdam, Potsdam, Germany

^22^Department of Agroecology, Aarhus University, Tjele, Denmark

^24^Swift Current Research and Development Centre, Agriculture and Agri-Food Canada, Swift Current, Saskatchewan, Canada

^25^ Soil Science Section, Faculty of Organic Agricultural Sciences, University of Kassel, Germany (before ^9^)

^26^Institute of Bio- and Geosciences - IBG-3, Agrosphere, Forschungszentrum Jülich GmbH, Jülich, Germany

**Variance components estimation of structure and parameter variance**

The R instructions that estimate structure, parameter and total variance are as follows:

lmerHere<-lmer(DAS~1+(1|structure),data=dataHere)

#The data frame dataHere has data for environment e, development stage s

\# DAS = days after sowing, structure = name of model structure.

lmerSig2Str<-attr(VarCorr(lmerHere)[[1]],"stddev")^2 # extract structure variance

lmerSig2Par<-sigma(lmerHere)^2 # extract parameter variance = variance of residuals

**Table S1**

**Model structures represented in this study.**

| Model structure | Version | References |
| --- | --- | --- |
| AgroC | May 2018 | (Herbst et al., 2008; Klosterhalfen et al., 2017) |
| APSIM | 7.8, 7.9, 7.10 | (Holzworth et al., 2014; Keating et al., 2003) |
| AquaCrop | 4.0 | (Vanuytrecht et al., 2014) |
| CERES-Wheat  with DSSAT | CERES in DSSAT V4.7. | (Hoogenboom et al., 2019b, 2019a) |
| CERES with Expert-N | Expert-N 5.0 | (Ritchie et al., 1988; Stenger et al., 1999) |
| CoupModel | Version 5.4.4 | (Coucheney et al., 2018; Jansson, 2012; Senapati et al., 2016) |
| CROPSIM-Wheat  with DSSAT | CROPSIM in DSSAT V4.7 | (Hoogenboom et al., 2019b; Hunt and Pararajasingham, 1995) |
| Cropsyst | 3.04.08 | (Stockle et al., 2001) |
| DAISY | 5.59 | (S. Hansen et al., 2012) |
| GECROS | Expert-N 5.0 | ( Stenger et al., 1999; Yin and van Laar, 2005) |
| HERMES | 4.27 | (Kersebaum, 2011, 2007) |
| LINTUL | LINTUL5 | (Wolf, 2012) |
| MONICA | 2.02 | (Nendel et al., 2011; Specka et al., 2019, 2015) |
| Nwheat | DSSAT | (Kassie et al., 2016) |
| OpenCrop |  | (Crout et al., 2018) |
| PANORAMIX | R version | (Chatelin et al., 2005; Gate, 1995) |
| Salus |  | (Basso et al., 2011; Basso and Ritchie, 2015) |
| SPASS with Expert-N | Expert-N 5.0 | (Stenger et al., 1999; Wang, 2000) |
| SSM-Wheat |  | (Soltani et al., 2013) |
| STICS | 8_5_0 | (Brisson et al., 2009; Coucheney et al., 2015) |
| SUCROS | Expert-N 3.0 | (Laar et al., 1992) |
| Wheat-Grow | 3.1 | (Lv et al., 2016; Zhu et al., n.d.) |
| WOFOST | 7.1.7 | (Boogaard et al., 1998) |

**Table S2**

**Parameter and structure variance averaged over environments of the French data sets. The variance components estimate is based on ensemble studies where some model structures were used by multiple modeling groups. The usual compared to protocol estimate is based on comparing usual and protocol calibration results for each modeling group. Parameter variance, structure variance and the ratio of parameter variance to total are averages over the values for each environment. The latter is different than the ratio of average parameter variance to average total variance.**

|  |  | Apache | | | Bermude | | | |
| --- | --- | --- | --- | --- | --- | --- | --- | --- |
| Method of estimating parameter variance as fraction of total | Development stage | BBCH10 | BBCH30 | BBCH55 | | BBCH10 | BBCH30 | BBCH55 |
| Variance components | Average number of modeling groups | 23 | 25 | 27 | | 23 | 25 | 27 |
|  | Number of model structures | 17 | 19 | 21 | | 17 | 19 | 21 |
|  | Parameter variance | 11.74 | 52.94 | 26.77 | | 12.05 | 71.25 | 36.41 |
|  | Structure variance | 5.05 | 4.29 | 9.65 | | 4.99 | 10.86 | 12.73 |
|  | Parameter variance as fraction of total | 0.55 | 0.93 | 0.81 | | 0.56 | 0.92 | 0.8 |
| Usual compared to protocol calibration | Average number of modeling groups | 14 | 16 | 17 | | 14 | 16 | 16.9 |
|  | Parameter variance | 0.75 | 16.62 | 10.35 | | 2.57 | 28.52 | 10.01 |
|  | Parameter variance as fraction of total | 0.28 | 0.88 | 0.71 | | 0.39 | 0.89 | 0.65 |

**Table S3**

**Parameter and structure variance averaged over environments of the Australian data set. The variance components estimate is based on ensemble studies where some model structures were used by multiple modeling groups. The usual compared to protocol estimate is based on comparing usual and protocol calibration results for each modeling group.**

| Method for estimating parameter variance as fraction of total | Development stage | BBCH10 | BBCH30 | BBCH65 | BBCH90 |
| --- | --- | --- | --- | --- | --- |
| Variance components | Average number of modeling groups | 22.95 | 23.93 | 27.64 | 26.05 |
|  | Average number of model structures | 17.95 | 19.93 | 22.64 | 21.05 |
|  | Parameter variance | 40.69 | 102.88 | 47.1 | 30.3 |
|  | Structure variance | 276.33 | 45.74 | 48.4 | 90.71 |
|  | Parameter variance as fraction of total | 0.12 | 0.66 | 0.49 | 0.24 |
| Usual compared to protocol calibration | Average Number of modeling groups | 13.9 | 16.9 | 18.9 | 17.3 |
|  | Parameter variance | 8.41 | 62.54 | 67.12 | 80.76 |
|  | Parameter variance as fraction of total | 0.026 | 0.65 | 0.61 | 0.46 |

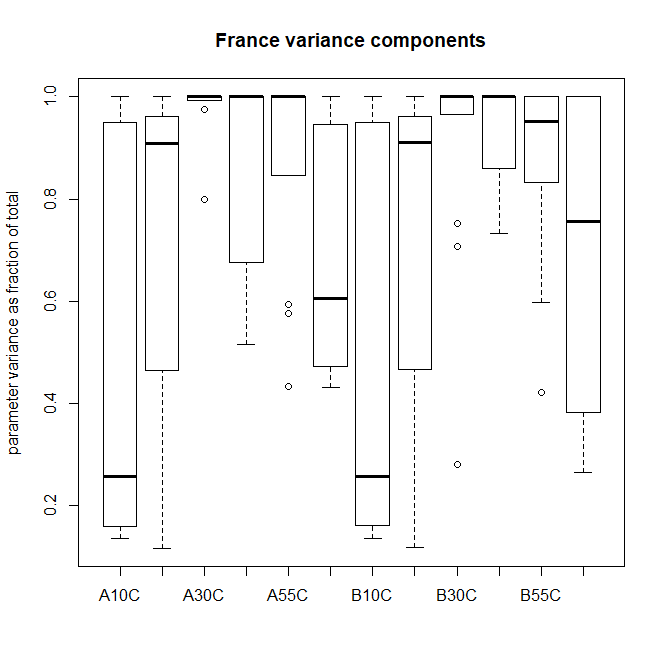

Fig S1. Boxplot of parameter variance for each environment as a fraction of total variance for the French data sets, based on variance components estimation. In the x axis labels, A=Apache data set, B= Bermude data set. 10, 30 and 55 refer to development stages BBCH10, BBCH30 and BBCH55 respectively. C indicates the calibration data subset. The boxplots without labels are for the same variety and development stage as the preceding boxplot, but for the evaluation data subset.

Fig S2. Boxplot of parameter variance for each environment as a fraction of total variance for the Australian data set, based on variance components estimation. In the x axis labels, 10, 30, 65 and 90 refer to development stages BBCH10, BBCH30, BBCH65 and BBCH90 respectively. E and C indicate the evaluation and calibration data subsets respectively.

Zhu, Y., Liu, L., Liu, B., n.d. WheatGrow: A simulation model for predicting growth andproductivity in wheat, in: The Workshop on Modeling Wheat Response to High Temperature. Texcoco, Mexico.
